# Deep Homology and Developmental Constraint Underlies the Replicated Evolution of Grass Awns

**DOI:** 10.1101/2024.05.30.596325

**Authors:** Erin Patterson, Dana MacGregor, Michelle Heeney, Joseph Gallagher, Devin O’Connor, Benedikt Nuesslein, Madelaine Bartlett

## Abstract

Replicated trait evolution can provide insights into the mechanisms underlying the evolution of biodiversity. One example of replicated evolution is the awn, an organ elaboration in grass inflorescences. Awns are likely homologous to leaf blades. We hypothesized that awns have evolved repeatedly because a conserved leaf blade developmental program is continuously activated and suppressed over the course of evolution, leading to the repeated emergence and loss of awns.

To evaluate predictions arising from our hypothesis, we used ancestral state estimations, comparative genetics, anatomy, and morphology to trace awn evolution. In line with our predictions, we discovered that awned lemmas that evolved independently share similarities in anatomy and developmental trajectory. In addition, in two species with independently derived awns and differing awn morphologies (*Brachypodium distachyon* and *Alopecurus myosuroides)*, we found that orthologs of the YABBY transcription factor gene *DROOPING LEAF* are required for awn initiation. Our analyses of awn development in *Brachypodium distachyon, Alopecurus myosuroides*, and *Holcus lanatus* also revealed that differences in the relative expansion of awned lemma compartments can explain diversity in awn morphology at maturity. Our results show that developmental conservation can underlie replicated evolution, and can potentiate the evolution of morphological diversity.

## Introduction

Phenotypic variation creates opportunities for natural selection to act in the evolution of biodiversity. Conserved developmental pathways can shape the character of this phenotypic variation. This shaping is usually viewed as a negative constraint, where developmental conservation limits phenotypic variation (Prusinkiewicz *et al*., 2007; Wessinger & Hileman, 2016; Blount *et al*., 2018). However, there are cases where development acts to generate or potentiate phenotypic variation (Gould, 2002; West-Eberhard, 2003; Rajakumar *et al*., 2012; Leichty & Sinha, 2021).

For example, ants in the genus *Pheidole* share ancestral developmental potential to generate a supersoldier caste. Supersoldier ants appear naturally in two species, but can be induced by hormones in three additional species, indicating a conserved developmental pathway with the potential to facilitate phenotypic change (Rajakumar *et al*., 2012). Here, we show that developmental conservation in the grasses can act to potentiate (rather than limit) morphological diversity. We hypothesize that developmental conservation underlies the replicated evolution of a morphological trait (James *et al*., 2023) - the awned grass lemma.

Lemmas are bract-like organs that subtend grass flowers (Sajo *et al*., 2008; Kellogg, 2015; Patterson *et al*., 2023). Many lemmas bear bristle-like extensions called awns (Fig. 1a,c). Awned lemmas have evolved multiple times, and lemma morphology and function varies extensively across the grasses (Linder *et al*., 2018; McAllister *et al*., 2018; Cavanagh *et al*., 2020; Petersen & Kellogg, 2022). Many lemmas have simple, short awns with no hypothesized functions (Ntakirutimana & Xie, 2019). However, longer and more complex awns can contribute to grass grain development, dispersal, and seedling establishment (Yanez *et al*., 2018; Cavanagh *et al*., 2020; Sanchez-Bragado *et al*., 2023); features that have likely contributed to the grass family’s extraordinary ecological success and dominance (Linder *et al*., 2018).

**Figure 1.**
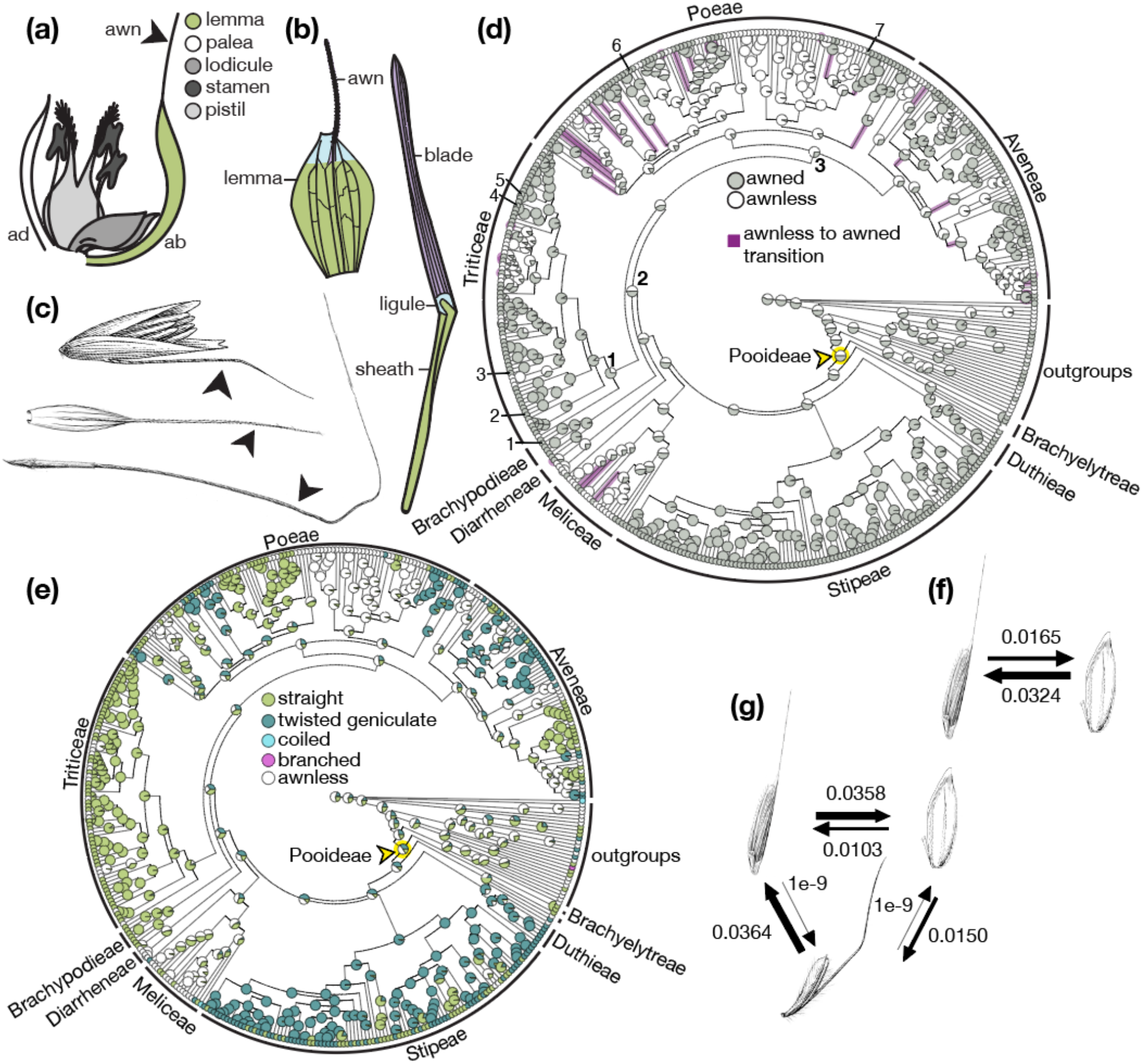
Awn presence and morphology have complex evolutionary histories in the Pooideae. (a) Generalized grass flower diagram, awn noted with arrow. (b) Predicted homology of awned lemmas to leaves. (c) Diversity of awns in the Pooideae. From top: abaxially insterted twisted geniculate awn (*Calamagrostis sp*.), apically inserted straight awn (*Secale cereale*), and apically inserted twisted geniculate awn (*Stipa spartea*). (d) Ancestral state reconstruction for awn presence. Species of interest noted: (1) brachypodium, (2) *Bromus tectorum*, (3) barley, (4) rye, 5) wheat, 6) velvetgrass, 7) blackgrass. Nodes of interest noted: most recent common ancestor of (1) Triticeae, (2) Brachypodium and Triticeae, (3) Poeae and Aveneae. Purple branches indicate gain of awns. Yellow node indicates Pooideae most recent common ancestor. e) Ancestral state reconstruction for awn morphology. f) Transition rates between awned and awnless states in (d). g) Transition rates between straight, twisted geniculate, and awnless in (e). Lemma and leaf in (b) redrawn from Thi-Tuyet-Hoa (1965). Images in (c), (f), and (g) from Hitchcock-Chase Collection of Grass Drawings.

Awn function is tied to awn form. For example, lemmas with barbed awns may function in animal fruit dispersal (Elbaum *et al*., 2007; Hua *et al*., 2015), while feathery awns may function in wind dispersal (Hensen & Müller, 1997; Yanez *et al*., 2018). Twisted geniculate awns, which have a proximal twisted column and knee-like bend, are often hygroscopic, and repeatedly twist to move grass grains across a surface (Peart, 1979; Raju, 2011). This movement may increase grain dispersal distance and help with seedling establishment by burying grains deeper underground (Garnier & Dajoz, 2001; Cavanagh *et al*., 2020; Morris, 2021). Awns in barley and wheat photosynthesize, and contribute photosynthate to developing grains (Abebe *et al*., 2009; Sanchez-Bragado *et al*., 2023). Therefore, awned lines of wheat and barley have fewer, heavier grains (Weyhrich *et al*., 1994; Motzo & Giunta, 2002; Sanchez-Bragado *et al*., 2023). Extra provisions in these heavier grains may help with seedling establishment (Nik *et al*., 2011; Linder *et al*., 2018; Muhsin *et al*., 2021; Petersen & Kellogg, 2022). Awns have also been implicated in positioning dispersed grains in the air column (during wind dispersal) and on soil (Peart, 1981), and in preventing herbivory (Schöning *et al*., 2004; Ceradini & Chalfoun, 2017; Titulaer *et al*., 2018), as well as modifying canopy temperature and shattering (Ntakirutimana & Xie, 2020). Thus, awned lemma forms and functions have diverged extensively.

Despite morphological and functional variation, all lemmas are leaf homologs (Thi-Tuyet-Hoa, 1965; Von Goethe & Miller, 2009; Patterson *et al*., 2023). Grass leaves contain three compartments along the proximal-distal axis - the sheath, the ligule, and the blade (Lewis & Hake, 2016). In awned lemmas, awns are likely homologous to the blade compartment ((Duval-Jouve, 1871; Thi-Tuyet-Hoa, 1965), Fig. 1b). Therefore, unawned lemmas are predicted to have lost blade compartments.

Morphological trait loss is often associated with parallel genetic loss. However, genetic loss is unlikely to be the case in awn evolution. Gene loss accompanied the evolution of eyelessness in cavefish (Sifuentes-Romero *et al*., 2020), of rootlessness in aquatic plants (Hepler *et al*., 2020; Ware *et al*., 2023), and of fungal cells that cannot crawl (Torruella *et al*., 2015; Prostak *et al*., 2021). Unlike these cases, awn loss is easily reversible: awns have been gained, lost, and regained many times over the course of grass evolution (Humphreys *et al*., 2011; Petersen & Kellogg, 2022). In addition, many grass species have unawned lemmas, but vegetative leaves with a typical leaf blade, including *Zea mays* (corn), and many *Oryza sativa* (rice), *Sorghum bicolor* (sorghum), and *Triticum aestivum* (wheat) cultivars (Kellogg, 2015). These data indicate that even when awns are not present, the genes that regulate leaf blade (and awn) development are maintained in grass genomes.

The retention of a blade developmental program in awnless species may provide a cache of genetic material that, under the right circumstances, could be activated to generate phenotypic variation upon which selection could act. We hypothesize that awns are gained and lost through the repeated activation and suppression of a leaf blade developmental program in lemmas. Under this hypothesis, developmental constraint would not limit phenotypic variation, but instead potentiate its emergence (Gould, 2002; West-Eberhard, 2003).

To test this hypothesis, we dissected awn evolutionary history in the grasses, focusing on the Pooideae subfamily, which includes the genetic experimental system *Brachypodium distachyon*, as well as many crops like barley, oats, and wheat (Schubert *et al*., 2019; Zhang *et al*., 2022). We evaluated three predictions arising from our hypothesis. Specifically, if a conserved developmental program underlies the replicated evolution of awns, we predicted that independently derived awns would have (1) conserved vascular anatomy; (2) conserved developmental trajectories; and (3) conserved genes regulating development. Our comparative analyses of anatomy, morphology, development, and genetics revealed the conserved developmental mechanisms underlying both the replicated evolution of awns, and the evolution of awn morphological diversity.

## Materials & Methods

### Ancestral State Estimation

We collected awn traits from GrassBase for species present in a time-calibrated phylogeny (Clayton *et al*., 2006; Schubert *et al*., 2019), and mapped the traits to the tree. We estimated ancestral states for each trait and rates of transition between traits using a maximum likelihood approach from corHMM (Boyko & Beaulieu, 2021). For each categorical trait, we compared two Markov models: Equal Rates (ER), which holds all transition rates between traits to be equal, and All Rates Different (ARD), where all transition rates may freely differ. In all cases, the ARD model was preferred (with a smaller AICc value), and was used with stochastic character mapping to plot the likely state at each ancestral node (Revell, 2012). Briefly, once the model has been generated, stochastic mapping uses a Bayesian approach to repeatedly generate random histories that allow the tree to arrive at the extant tip states (Bollback, 2006). These repeated simulated histories were combined and the proportion of each state at each node was plotted. For phylogenetic signal of the binary presence trait, the *D*-statistic was calculated using the R package caper. A *D* value of 1 indicates a trait is distributed randomly across a phylogeny, whereas a *D* value of 0 indicates strong phylogenetic signal, consistent with a trait evolving under Brownian motion (Fritz & Purvis, 2010; Orme, 2013).

### Plant Materials & Growth Conditions

Brachypodium *awnless1* mutant grains were developed by fast neutron mutagenesis in the Bd21 background (Derbyshire & Byrne, 2013). Genotyping of *awl1* was performed via multiplex PCR with primers spanning the deletion and a pair specific to a gene inside the deleted region (Table S1).

Blackgrass and velvetgrass grains were obtained from the USDA-ARS GRIN (Germplasm Resources Information Network) or from germplasm held at Rothamsted Research. For anatomical analysis of blackgrass, accessions PI 289645 and PI 204402, originating from Spain and Turkey respectively, were used. For VIGS in blackgrass, a line derived from the Peldon biotype was used (Mellado-Sánchez *et al*., 2020). For velvetgrass, accessions PI 302907 and PI 311415, both from Spain, were used. Brachypodium and velvetgrass were grown under long day conditions (20h light, 4h dark), at 28°C. Blackgrass was grown under short day conditions (10h light, 14h dark), at 22°C except for VIGS, details for which are given below.

Awns were collected from various species as follows: wheat and rye growing at the UMass Crop Research Center, ‘Steptoe’ barley grown in the UMass Morrill Glasshouse, and *Bromus tectorum* (cheatgrass) growing outside the UMass Morrill Science Center.

### Staining & Microscopy

Awn and leaf sections were made by hand and immediately moved to sterile water on a microscope slide to keep fresh. When several sections had been collected, the water was exchanged for Toluidine Blue for 1-2 minutes, after which the sections were washed to remove excess stain. A coverslip was applied, and sections were imaged. For stained sections, a Zeiss Axioplan microscope with AmScope digital camera (MU633-BI) was used.

For scanning electron microscopy, a JEOL JCM-6000Plus was used. For developing flowers, fresh tissue was dissected immediately before imaging and mounted on a stub with oven-bake clay (Sculpey III). For senesced awns, the whole awn was mounted in the same manner. In all cases, images were collected under high vacuum and low voltage (5kV).

### Awn Clipping

The awn clipping experiment was performed twice, in the same manner except where noted. Brachypodium grains from the Bd21-3 inbred line were grown in 2.5 inch pots, with four germinants per pot. In each pot, plants were randomly assigned to a treatment, with one glume removal, one no treatment, and two awn removal plants. At flowering, in the awn removal treatment, awns were cut off as they appeared. In the glume removal treatment, the lower glume on each spikelet was cut off on the first day that awns were visible. Plants in the no treatment condition were not clipped, but were handled a similar amount as the other two treatments.

In the second replicate experiment, at senescence, all spikelets were collected, and the number of filled grains per spikelet was counted (light blue, Fig. 2d). The first replicate was the same, but only collected spikelets with one or more filled grains (dark blue, Fig. 2d). Filled grains were weighed. To control for differences in weight due to the absent awn, two different methods were used. In the first replicate (dark blue, Fig. 2c), the lemmas were fully removed from all grains before weighing. In the second replicate (light blue, Fig. 2c), the awns were cut off of the glume and no treatment lemmas before weighing. Due to their small size, grains were weighed while grouped by spikelet, and the total weight was divided by the number of grains.

**Figure 2.**
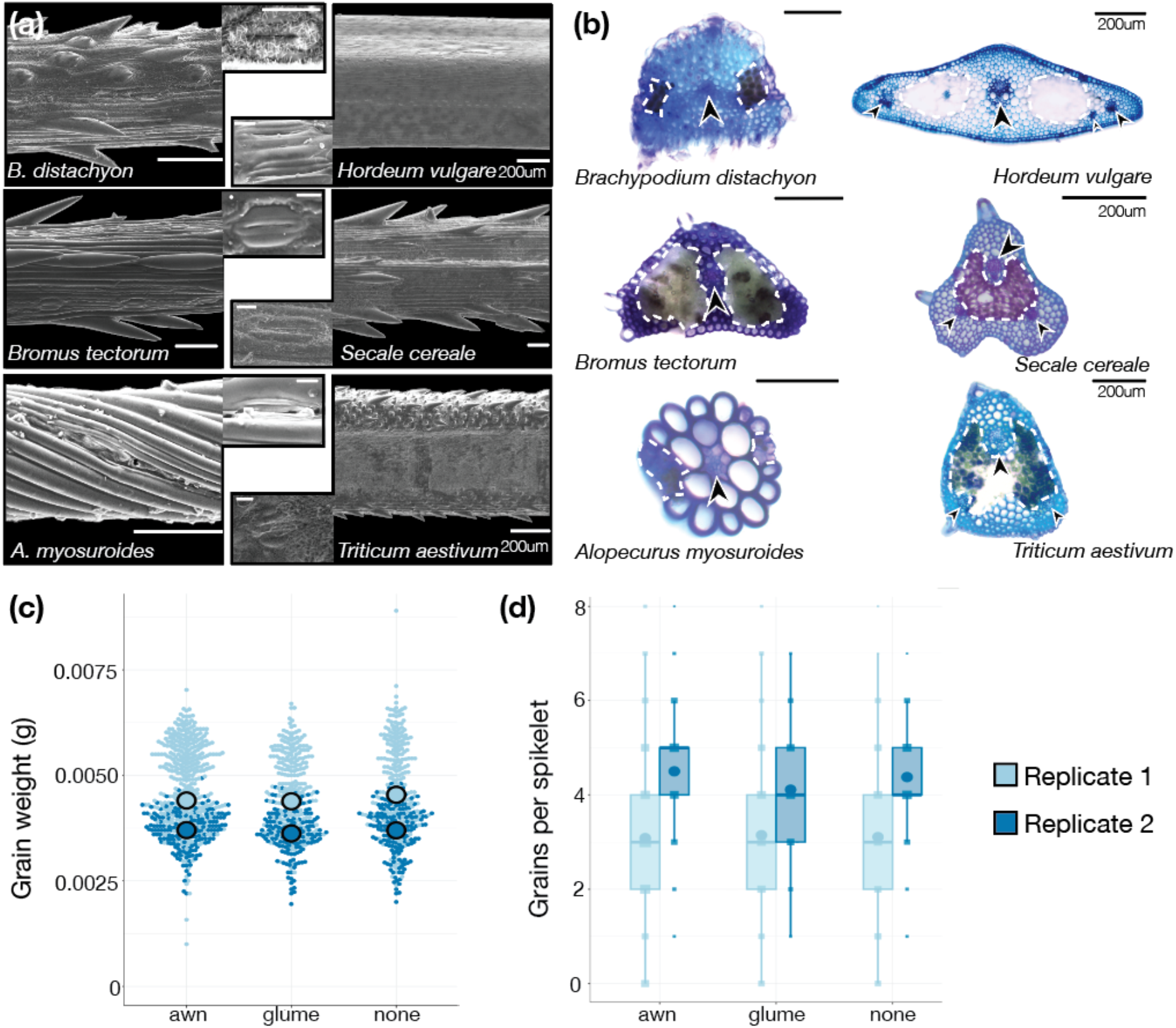
Structure, but not function, of awns is conserved. (a) Awns from six Pooideae species, stomata in insets (10uM scale bars). (b) Transverse sections of awns from the same six species, chlorenchyma (white dotted lines) and vasculature (arrows) noted. Scale bars 100um throughout unless noted. (c) Awn removal has no effect on grain weight in brachypodium. (d) Awn removal has no effect on filled grains per spikelet in brachypodium. Light and dark blue denote data from replicate experiments.

### Genomic data analysis

Illumina 150 bp paired end reads from *awl1* were trimmed and mapped to the *Brachypodium distachyon Bd21* genome using GSNAP (International Brachypodium Initiative, 2010; Bolger *et al*., 2014; Wu *et al*., 2016). Because *awl1* was the product of fast neutron mutagenesis, we expected that a large deletion would be the most likely mutation. Paired end reads that mapped uniquely but at a much greater distance than expected (‘PL’ flag in GSNAP) were visually inspected in IGV to find deletions (Thorvaldsdóttir *et al*., 2013).

*CRISPR-Cas9 Gene Editing in Brachypodium* Spacers targeting *BdDL* (Bradi1g69900) were designed using CRISPOR (Concordet & Haeussler, 2018). These spacers were synthesized and assembled into a guide RNA construct using the MoClo system (Miao *et al*., 2013; Engler *et al*., 2014). An additional MoClo construct was assembled containing the ubiquitin promoter from maize driving a Cas9 with introns (Grützner *et al*., 2021). These constructs were combined and Gateway cloned into a hygromycin-containing resistance vector (O’Connor *et al*., 2014). This construct was introduced to brachypodium Bd21-3 embryogenic callus via *Agrobacterium-*mediated transformation using strain AGL1 (Vogel & Hill, 2008). Calli were screened with hygromycin resistance, and *Bddl-CR* alleles were genotyped via PCR and Sanger sequencing.

### RT-qPCR

Inflorescence meristems were collected and pooled from three WT and three *awl1* plants, and immediately frozen. Total RNA was extracted using the Qiagen RNeasy Plant Kit, following the kit instructions. 500ng of RNA was used for cDNA synthesis, using the SuperScript III RT kit (ThermoFisher Scientific). qPCR was performed with primers specific to BdDL, as well as control primers for BdUBC18 (Hong *et al*., 2008).

### VIGS in Blackgrass

VIGS was performed in blackgrass following established protocols (Lee *et al*., 2015; Mellado-Sánchez *et al*., 2020). Briefly, two fragments targeting regions in the gene of interest (*AmDL*) were cloned in antisense into modified Barley Stripe Mosaic Virus (BSMV) constructs following protocols described in Lee et al. (Lee *et al*., 2015). Two control vectors carrying the empty multiple cloning site (‘MCS’) from Mellado-Sanchez et al. (2020), or a 230bp region from *GFP* (GenBank E17099.1) targeting from 307 to 536 from the start site made for this project were also used. A vector targeting an antisense portion of *PHYTOENE DESATURASE* (*AmPDS*) from Mellado-Sanchez et al. (2020) was also used as a visual positive control to check efficiency of VIGS. These showed clear photobleaching at 96.5% efficiency two weeks after inoculation. Sap extracted from *N. benthamiana* leaves infected with the different BSMV constructs was used to inoculate young black-grass seedlings two weeks after they had been vernalized for five weeks at 5°C. The plants were grown at 27°C with 16 hours of light and 21°C with 8 hours of dark for two weeks after rub inoculation to allow the virus to establish and then moved to 21°C with 16 hours of light and 15°C with 8 hours of dark until flowering.

## Results

### Awns have been gained, lost, and re-gained many times in the Pooideae

To examine the evolutionary history of awns, and to identify cases of independent awn emergence for further study, we reconstructed the most likely ancestral awn states in the Pooideae, focusing on lemma awns (Fig. 1c). We mapped awn presence onto a time-calibrated tree containing about 10% of the species in the subfamily (Schubert et al., 2019) (Fig 1d). Reconstructions at deep nodes were ambiguous, with both awned and unawned lemmas predicted to be equally likely in the common ancestor of all Pooideae species. However, we predicted several instances of likely gain and loss of awns at shallower nodes. Awns were gained roughly twice as often as they were lost in the Pooideae, with awns having been gained at least 25 times independently, and lost at least 14 times in our sampling (Fig. 1d,f). These numbers are likely an underestimate, given that the phylogeny we used includes only a small fraction of Pooideae species. We also estimated the phylogenetic signal of awn presence using the *D*-statistic (Fritz & Purvis, 2010). Phylogenetic signal provides an estimate of the tendency of a trait to be similar between closely related species, rather than evolving randomly across a phylogeny (Kamilar & Cooper, 2013). The *D*-statistic was estimated at 0.2667, indicating that awn presence was not evolving randomly (p = 0). Thus, awn evolution is a labile but not random process in the Pooideae. To assess how awn morphology changed across the Pooideae, we mapped awn morphology, awn length, and awn insertion point (Fig. 1e, Fig. S1,S2). Most species in our dataset showed one of three morphologies: (1) no awns, (2) twisted geniculate awns, or (3) straight awns (Fig. 1c). Coiled or branched awns were present only in *Streblochaete longiarista* and in some outgroup species. As with awn presence, character states at deeper nodes were ambiguous, but shallower nodes within tribes showed clear patterns. For example, the Triticeae (wheat, barley, and relatives) only had straight awns. In contrast, the other large tribes (Poeae, Aveneae, and Stipeae) all contained a mix of species with each awn type. The longest awns were specific to Stipeae and wheat, barley, and their close relatives (Fig. S1), reaching up to 25 times the length of the lemma body, as in *Nassella tenuissima* (Stipeae). On average, awns in this subset of the Pooideae are about 3.6 times the length of the lemma, and only 31% of awned species have awns equal to or shorter than the length of their cognate lemma bodies. Awns were abaxially inserted in species from the Meliceae, Poeae, and Aveneae, as well as in a single Triticeae species (Fig. S2). Transition rates for morphology show that straight awns rarely shifted to twisted geniculate, but twisted geniculate often shifted to straight (Fig. 1g). Additionally, twisted geniculate awns have evolved from the awnless state about as often as straight awns have (Fig. 1g). Overall, our analyses indicate that awn morphology is diverse in the Pooideae. Both straight and twisted geniculate awns have been independently derived multiple times, suggesting these independently derived awns serve diverse functions in the Pooideae.

### Micromorphology varied, but vascular anatomy was similar between independently derived awns

Conserved vascular anatomy suggests homology between plant organs (Li & Xu, 2020; Hsu & Kuo, 2021). Therefore, if developmental constraint underlies the repeated evolution of awns, we would expect the vascular anatomy of independently derived awns to be conserved. To evaluate this prediction, we examined anatomy and micromorphology in six Pooideae species representing two to three independent awn derivations (Fig. 2a,b). First, awns in *Alopecurus myosuroides* (blackgrass) were independently derived from those in the other five species we examined (Fig. 1d, node 3). Second, the Triticeae species *Triticum aestivum* (wheat), *Hordeum vulgare* (barley), *Secale cereale* (rye), and *Bromus tectorum* (cheatgrass) likely share a derivation (Fig. 1d, node 1). Third, awns in brachypodium may have been independently derived from those in the Triticeae, but this prediction was more ambiguous (Fig. 1d, node 2). Blackgrass has twisted geniculate awns, while wheat, barley, rye, cheatgrass, and brachypodium all have straight awns.

Micromorphology varied, but vascular anatomy was similar between the surveyed species. All six species had stomata in rows along the abaxial surface of the awn. Both presence and characteristics of other micromorphological traits varied between species, including cuticular wax formation, trichome presence and density, and trichome types (Fig. 2a). In cross-section, the straight awns all had many small cells, while the twisted geniculate awn of blackgrass was composed of few large cells, with only two cell layers surrounding the central vascular bundle (Fig. 2b). In contrast, all surveyed awns had a central vascular bundle and two regions of lateral chlorenchyma (Fig. 2b). Together, these results suggest that micromorphology is not conserved (e.g., presence of hair cells), while vascular anatomy is largely conserved across independently derived awns. Variation in micromorphology suggests divergent functions (e.g. in herbivore deterrence), while conserved vascular anatomy suggests shared evolutionary origins (Song *et al*., 2022).

### Awn removal in brachypodium did not affect grain mass

We next examined whether function is conserved between the awns of brachypodium and wheat or barley. These species all have straight awns with similar cell types and organization. However, the awns of brachypodium are much shorter and have a smaller proportion of chlorenchymous tissue, and may have been independently derived from those in wheat and barley (Fig. 2b, Fig. S1,S2). Across many independent investigations, the awns in wheat and barley consistently contribute to grain mass, likely by contributing photosynthate to developing grains (Harlan & Hulton, 1920; Miller *et al*., 1944; Grundbacher, 1963; Li *et al*., 2006, 2023; Abebe *et al*., 2009; Maydup *et al*., 2010; Liller *et al*., 2017; Swarts *et al*., 2017; Sanchez-Bragado *et al*., 2023). To determine whether this was the case in brachypodium, we performed an awn removal experiment. In two independent experiments, we either (1) removed all awns, or (2) a single glume from developing spikelets (to control for wounding), or (3) left plants untreated. grain mass and the proportion of grains filled was similar across all three conditions (Fig. 2c,d).

Thus, despite similar construction (Fig. 2b), awn removal in brachypodium did not affect grain mass, as it does in barley and wheat (Harlan & Hulton, 1920; Miller *et al*., 1944; Grundbacher, 1963; Li *et al*., 2006, 2023; Maydup *et al*., 2010;

Liller *et al*., 2017; Swarts *et al*., 2017; Sanchez-Bragado *et al*., 2023).

### Independently derived awns had similar developmental trajectories

If a latent leaf developmental program underlies the repeated evolution of awned lemmas, then we expect awned lemma development to follow the stereotypical grass leaf developmental pattern. In grass leaves, the blade and sheath compartments both initiate early, but the blade begins expanding first. The sheath compartment expands later, concomitantly with the ligule (Lewis & Hake, 2016; Strable & Nelissen, 2021). To evaluate whether awned lemmas followed this pattern, we tracked awn development in three species with independently derived awns (Fig. 1c) and differing awn morphologies (Fig. 3): brachypodium (straight), blackgrass (abaxially-inserted twisted geniculate), and *Holcus lanatus* (velvetgrass, abaxially-inserted nearer the tip of the lemma, twisted geniculate). Velvetgrass was not included in anatomical analyses due to the small diameter (<100 um) and length (∼2 mm) of its awns, and limited availability of flowers.

**Figure 3.**
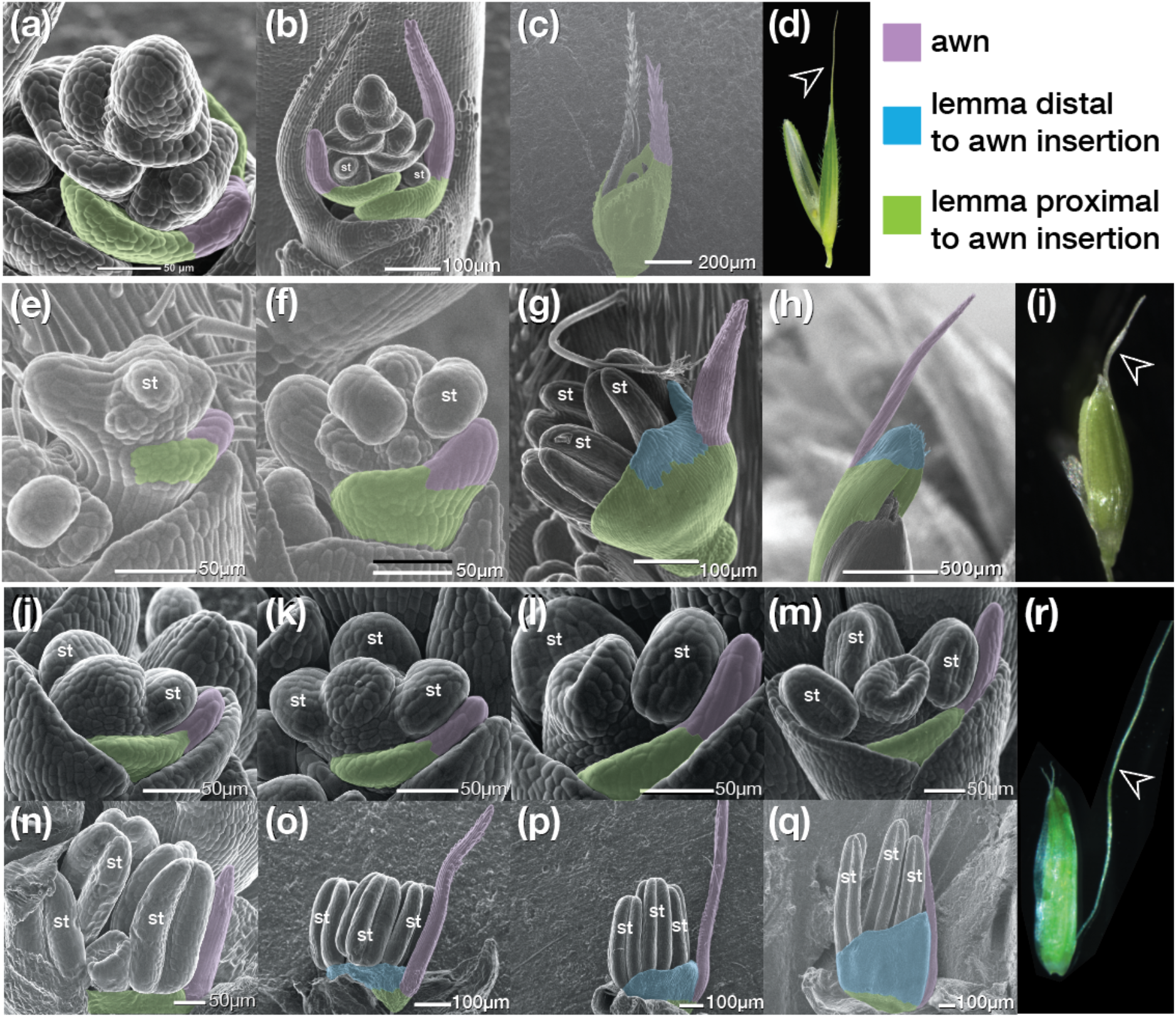
Developmental trajectory of awns is shared across three species with independently derived awns. (a-c) Scanning electron micrographs of awn development in brachypodium. (d) Mature brachypodium flower, awn noted with arrow. (e-h) SEMs of awn development in velvetgrass. (i) Mature velvetgrass flower, awn noted with arrow. (j-q) SEMs of awn development in blackgrass. (r) Mature blackgrass flower, awn noted with arrow. (s) Predicted homology of awned lemmas to leaves. Throughout, SEMs are false colored to indicate putative leaf compartment homologies. st = Stamen or stamen primordium expands later, concomitantly with the ligule (Lewis & Hake, 2016; Strable & Nelissen, 2021). To evaluate whether awned lemmas followed this pattern, we tracked awn development in three species with independently derived awns (Fig. 1c) and differing awn morphologies (Fig. 3): brachypodium (straight), blackgrass (abaxially-inserted twisted geniculate), and *Holcus lanatus* (velvetgrass, abaxially-inserted nearer the tip of the lemma, twisted geniculate). Velvetgrass was not included in anatomical analyses due to the small diameter (<100 um) and length (∼2 mm) of its awns, and limited availability of flowers.

In brachypodium, the lemma primordium appeared very early, subtending the floral meristem, with an acute tip. This acute tip extended into an awn with each subsequent, younger awn slightly shorter than the older one on the lemma below it (Fig. 3a,b). Once the stamens had begun to differentiate, the awn differentiated at a similar pace, producing hairs from tip to base (Fig. 3b). The sheath-like lemma compartment extended relatively late in development, once the carpels had begun to differentiate (Fig. 3c). Brachypodium lemmas had no clear ligule-like compartment.

In velvetgrass (*Holcus lanatus*), the lemma primordium also appeared early and had an acute tip (Fig. 3e). Unlike brachypodium, the velvetgrass spikelet only had one awned flower. The awned lemma expanded while the floral organs developed, with both the acute tip and lower lemma tissue expanding (Fig. 3f). Later, the awn extended and produced hairs as the stamens continued differentiating, and the ligule-like compartment distal to the awn insertion point extended (Fig. 3g). Finally, the sheath-like compartment proximal to the awn insertion point (the lemma body) extended as the floral organs matured (Fig. 3h).

In blackgrass, the lemma primordium appeared early in floral development, and had an acute tip, as in brachypodium and velvetgrass (Fig. 3j). However, in contrast to brachypodium and velvetgrass, the lemma then paused in development, while the inner floral organs initiated and matured (Fig. 3k, l). Only when the anthers had begun forming locules did the awn extend (Fig. 3m). Once the awn extended, the ligule-like lemma tissue distal to the awn insertion point appeared and extended (Fig. 3o-q). At maturity, the awn was inserted very near the base of the lemma, and the sheath-like compartment extended minimally (Fig. 3r). Thus, the majority of the lemma body in blackgrass is likely homologous to the ligule, rather than to the sheath, again in contrast to brachypodium and velvetgrass.

Despite differences in morphology, these species share a trajectory of (1) initiation of the sheath-like lemma base and blade-like awn; (2), extension and differentiation of blade-like awn; (3) simultaneous extension of the ligule-like distal lemma tissue (if present) and the sheath-like compartment. This trajectory is similar to the pattern of grass leaf development (Lewis & Hake, 2016; Strable & Nelissen, 2021), and adds support to the hypothesis that the awn is homologous to the leaf blade (Duval-Jouve, 1871; Thi-Tuyet-Hoa, 1965). Importantly, this trajectory is shared between independently derived awns with differing morphologies, suggesting a similar developmental program acting in all three species.

### DROOPING LEAF was necessary for awn development in independently derived awns with differing morphologies

If developmental conservation underlies the replicated evolution of awns, then we expect the same genetic pathways to regulate the independent derivations of awns. To evaluate this prediction, we explored the function of the YABBY transcription factor gene *DROOPING LEAF (DL)*, which regulates awn development in rice and barley (Toriba & Hirano, 2014; Zhang *et al*., 2024), and carpel and leaf blade midrib development in rice and maize (Ohmori *et al*., 2011; Toriba & Hirano, 2014; Strable *et al*., 2017; Strable & Vollbrecht, 2019).

We first mapped and cloned an existing brachypodium mutant, *awnless1 (awl1*) that phenocopies other *dl* mutants (Yamaguchi *et al*., 2004; Derbyshire & Byrne, 2013; Strable & Vollbrecht, 2019). The *awl1* mutant is awnless and lacks leaf midribs (Fig. 4b,f). We identified a deletion upstream of *BdDL* which is associated with significantly decreased expression of *BdDL* in the *awl1* mutant (Fig. 4g,j). Importantly, this deletion co-segregates with the *awl1* phenotypes (p =0.718, chi-square test; Fig. S3). We also generated a *Bddl* mutant using CRISPR-Cas9 genome editing, which recapitulated the awnless phenotype (Fig. 4h). This mutant was biallelic, with single base pair edits in exons 2 and 4 predicted to result in premature stop codons (Fig. 4i). Thus, *DL* is necessary for awn development, specifically awn initiation, in brachypodium.

**Figure 4.**
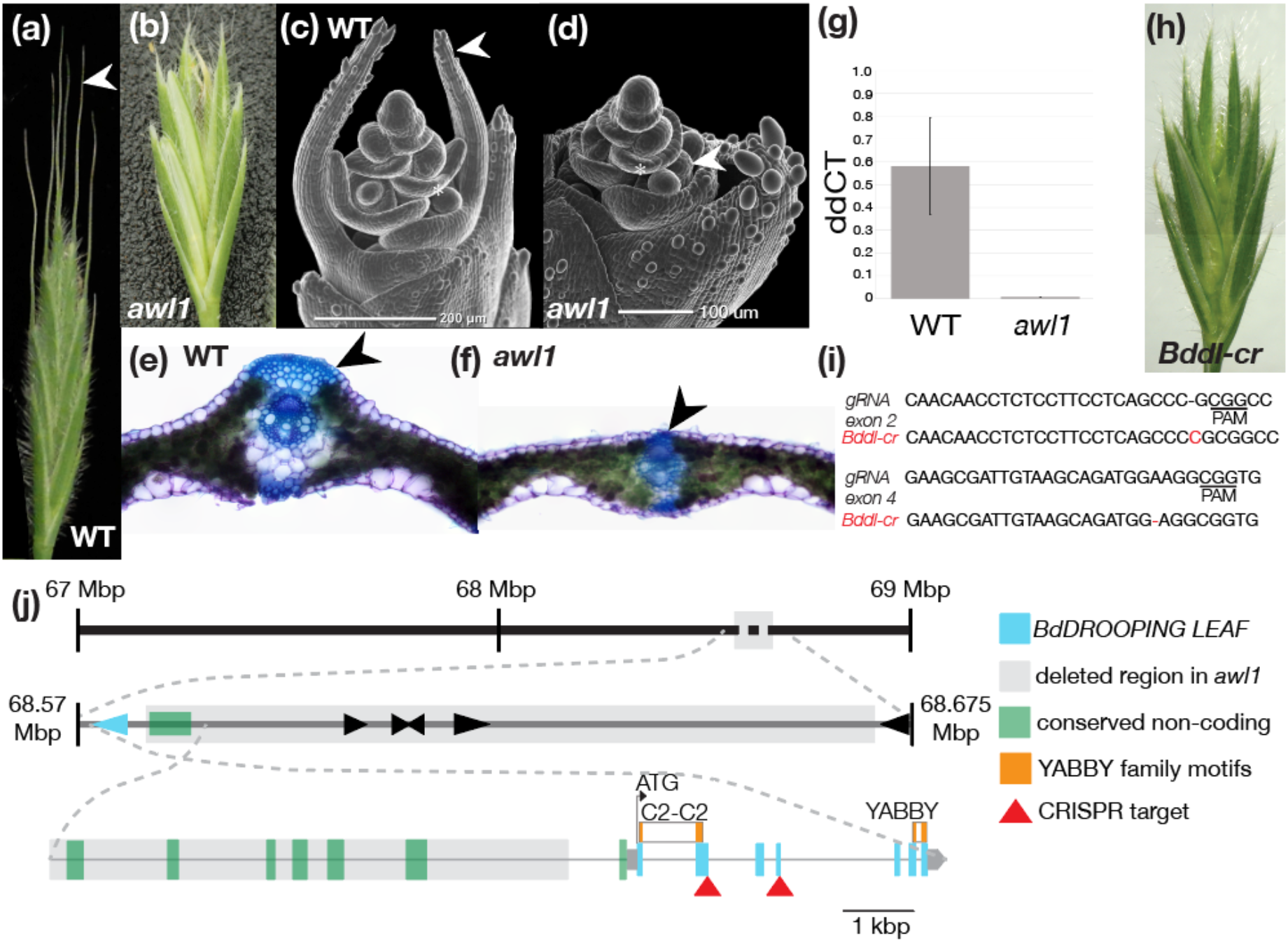
*BdDROOPING LEAF (BdDL)* is necessary for awn development in brachypodium. (a,b) Spikelet phenotypes (WT, *awl1* mutant), awns noted with arrow. (c,d) The *awl1* mutant does not initiate awn growth. Stamen primordia of similar age marked with asterisk. (e,f) The *awl1* mutant lacks leaf midribs (arrow). (g) The *awl1* mutant has very low expression of *BdDL* in floral tissue. (h) The *Bddl-cr* mutant phenocopies *awl1*, lacking awns. (i) Edited sequence in *Bddl-cr*, biallelic single nucleotide mutations in exons 2 and 4.(j) Genomic region of chromosome 1 of interest in the *awl1* mutant (top), with deletion (top, middle, and bottom, gray box), *BdDL* (middle and bottom, blue arrow and boxes (exons)), other genes (middle, black arrows), upstream conserved noncoding sequence (middle and bottom, green boxes). CRISPR construct targets shown (bottom, red arrows).

We next tested whether *DL* is necessary for awn development in blackgrass using virus-induced gene silencing (VIGS). The twisted geniculate awns in blackgrass were derived independently from the straight awns of brachypodium and rice (Fig. 1c). We used VIGS (Mellado-Sánchez *et al*., 2020) to knock down *AmDL* (ALOMY6G44958) expression in blackgrass. Blackgrass has a single *DL* homolog, reducing the likelihood of off-target effects (Cai *et al*., 2023). We designed three constructs targeting different regions of the *AmDL* mRNA, and a construct containing 230bp of *GFP* (GenBank: E17099.1) as a negative control for off-target effects.

Targeting *AmDL* with VIGS resulted in differences in awn initiation, length, and placement. Despite differences in efficiency, all three constructs targeting *AmDL* resulted in awnless lemmas (12/23 plants, total of 15 spikes, Fig. 5e). The two constructs with the strongest response also resulted in aberrant carpels, as in *dl* mutants in rice and maize (Nagasawa *et al*., 2003; Strable & Vollbrecht, 2019). All plants treated with the *GFP* or multiple cloning site (empty vector) negative control constructs had normal awn development (24/24 plants, Fig. 5a-c, Fig. S4).

**Figure 5.**
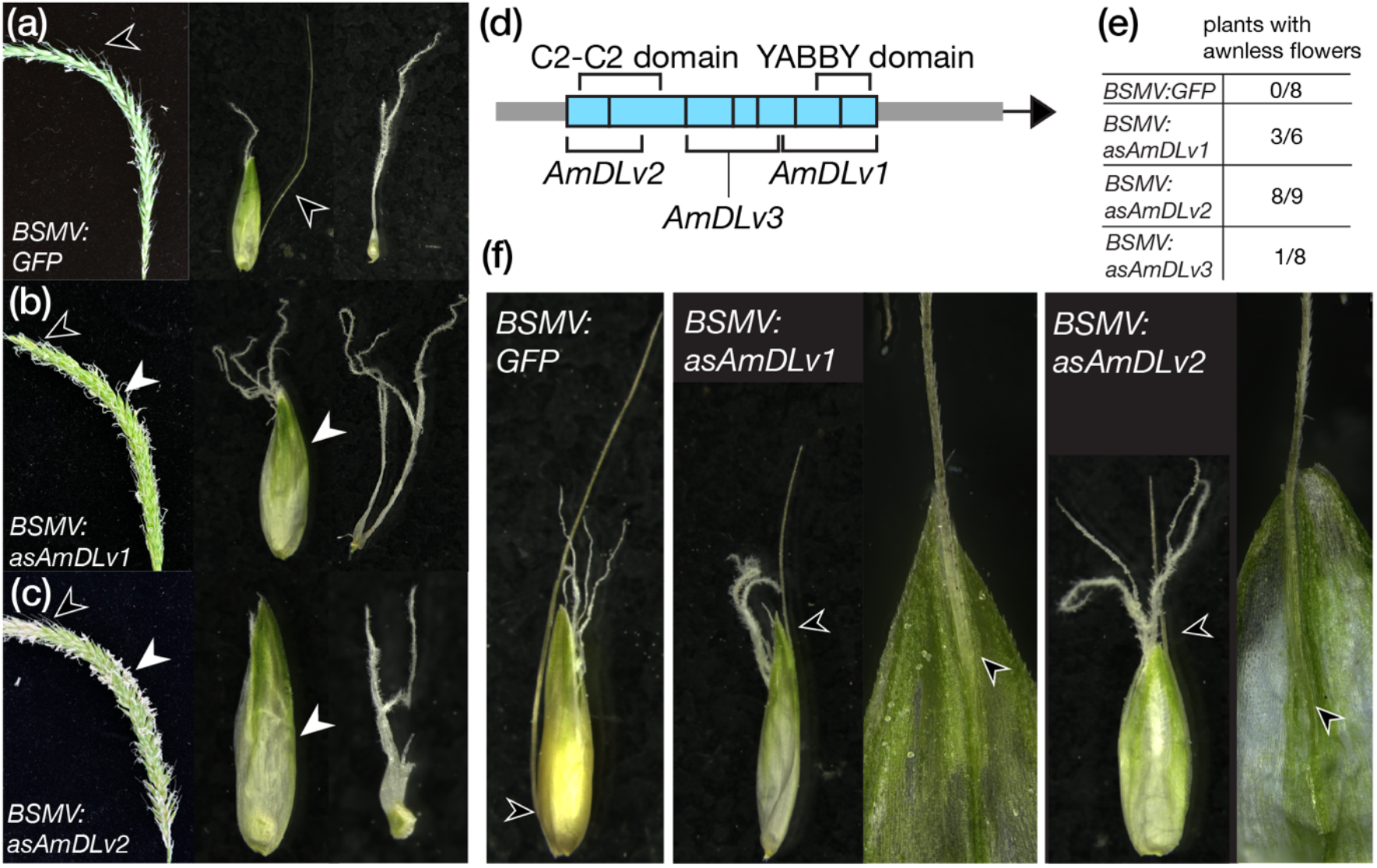
*AmDROOPING LEAF (AmDL)* is necessary for awn development in blackgrass, a species with twisted geniculate awns. (a) Spike, flower, and pistil from a representative plant from the *BSMV:GFP* control treatment, with awns present (black arrows). (b,c) Spike, flower, and pistil from the *BSMV:asAmDLv1* and *v2* treatments, with awns absent in center of spike (white arrows), and aberrant pistil morphology (right). (d) Regions of *AmDL* targeted by the *BSMV:asAmDL* constructs. (e) Number of plants with awnless flowers for the *GFP* control and three *AmDL* constructs. (f) Flowers from the *BSMV:asAmDLv1* and *v2* constructs with shorter awns also had awns inserted higher up on the lemma body.

Blackgrass awns are normally inserted near the base of the lemma. Interestingly, in more than half (8/15) of the *AmDL-VIGS* spikes with awnless flowers, other flowers on the affected spike had shorter awns. In these flowers, the awn was inserted near the center of the lemma and partially fused to the lemma, rather than free as in untreated and control blackgrass plants (Fig. 5e). This change in insertion point suggests that *DL* has a role in partitioning the leaf primordium along the proximal-distal axis. This could provide a mechanism for modulating awn insertion point, often an important taxonomic trait (Kellogg, 2015; Peterson *et al*., 2019). Together, these results show that *DL* is necessary for awn development in at least three independent derivations of awns (Fig. 1c) - rice (Toriba & Hirano, 2014), brachypodium (Fig. 4b), and blackgrass (Fig. 5b,c). A *DL* homolog also regulates awn development in barley (Zhang *et al*., 2024), which may represent a fourth independent derivation of awns (ancestral state estimations equivocal, Fig. 1c). Thus, although awns in these species are independently derived and differ in form and function, *DL* is a key regulator of awn development in all cases.

## Discussion

Despite a complex evolutionary history with many independent derivations, awns are likely homologous to leaf blades, and share developmental patterns and pathways. Awns evolved independently many times in the Pooideae (Fig. 1c) (Humphreys *et al*., 2011; Teisher *et al*., 2017; Zhang *et al*., 2022; Petersen & Kellogg, 2022). Vascular anatomy was similar between brachypodium, wheat, and barley awns (Fig. 2b), but awn function was not conserved; awns in brachypodium had no effect on measured grain traits in our experimental conditions (Fig. 2c). Independently derived awns, with differing morphologies and hypothesized functions, had developmental trajectories similar to each other and to grass leaves (Fig. 3) (Lewis & Hake, 2016; Strable & Nelissen, 2021). Further, *DL* was necessary for awn initiation in both brachypodium and blackgrass, and may have a role in regulating awn insertion point (Fig. 4, 5). Together, these data matched our predictions that independently derived awns have (1) conserved vascular anatomy; (2) conserved developmental trajectories; and (3) conserved genes regulating their development. Thus, a conserved leaf blade developmental program may underlie the replicated evolution of awns.

Many other examples of replicated evolution are clearly adaptations to particular environmental conditions: root loss in aquatic Lemnaceae, xerophytic forms in *Euphorbia*, eye loss in cavefish, benthic phenotypes in sticklebacks, and mandible shape in beetles (Horn *et al*., 2012; McGee & Wainwright, 2013; Baulechner *et al*., 2020; Sifuentes-Romero *et al*., 2020; Ware *et al*., 2023). For example, in beetles, mandible shape is strongly correlated with diet, regardless of phylogenetic history (Baulechner *et al*., 2020). While awns are likely homologous, they differ in morphology and hypothesized function, and many awns may lack specific functions. These differences indicate that the replicated evolution of awns is not always adaptive. Instead, awns may arise as frequently as they do as a consequence of development, and provide phenotypic variation for natural selection to act on in some circumstances.

Diversity in awned lemma morphology may be driven, in part, by differences in relative growth of lemma compartments. In the three species we examined closely, the expansion of the ligule-like compartment dictated awn insertion point. In brachypodium, no ligule-like compartment was visible, leading to a long apically inserted awn (Fig. 3d). In velvetgrass, there was a small ligule-like compartment, leading to a short sub-apically inserted awn (Fig. 3i). In blackgrass, the ligule-like compartment expanded far more than the sheath-like compartment. Therefore, the majority of the lemma body comprised ligule-like tissue, leading to a long, abaxially inserted awn (Fig. 3r). This indicates that different relative expansion of the three compartments (sheath-like, ligule-like, blade-like) was an important factor driving morphological differences between awned lemmas in these three species. This is similar to differential growth in leaves, generating morphological variety (Kaplan, 2001), or in heteromorphic flowers (Webster & Gilmartin, 2006).

Many aerial organs, including lemmas and floral organs such as carpels, are modified leaves (Von Goethe & Miller, 2009). Therefore, developmental programs may be common to several types. In addition to its role in leaf and awn development, *DL* also regulates carpel development (Toriba & Hirano, 2014; Strable & Vollbrecht, 2019). The brachypodium *awl1* mutant, blackgrass treated with *BSMV:asAmDL*, maize mutants *Zmdrl1* and *Zmdrl2*, and rice *dl* mutants all have aberrant carpel development (Ohmori *et al*., 2011; Toriba & Hirano, 2014; Strable *et al*., 2017; Strable & Vollbrecht, 2019). Similarly, the *Arabidopsis thaliana DL* ortholog *CRABS CLAW* (*CRC*), regulates nectary and carpel development (Lee *et al*., 2005), although *CRC* may have an ancestral role in leaf development (Fourquin *et al*., 2014). *DL* is not alone in having dual roles in awns and carpels: a *SHORT INTERNODES/STYLISH (SHI)* homolog has a role in both awn and carpel development in barley (Yuo *et al*., 2012). In rice, an epidermal patterning factor-like (EPFL) protein regulates both awn length and grain length, which is likely to be associated with carpel size (Jin *et al*., 2016). A growing number of genes are implicated in both carpel and lemma development (Zhang *et al*., 2016; Ma *et al*., 2019; Shoesmith *et al*., 2021; Jing *et al*., 2023). In some of these cases, carpel development is not directly limited by gene function, but rather by physical constraint of the developing grain due to decreased lemma size (Ren *et al*., 2018; Zhang *et al*., 2024). For those where physical constraint was not a factor, we suggest that dual roles for genes in lemma and carpel development are due to their shared evolutionary origins (Von Goethe & Miller, 2009). Ultimately, both carpels and lemmas are modified leaves, and may use the same genetic pathways during development.

Developmental constraints are often considered only in a restrictive sense, preventing phenotypes from being accessed. We highlight the positive ability of developmental constraints to facilitate replicated evolution. Our results highlight the fundamental relatedness of genetic processes in plants, and suggest that positive developmental constraint may be prevalent in instances of replicated evolution in plants, where many aerial organs are homologous to leaves (Von Goethe & Miller, 2009).

## Acknowledgements

We thank Amanda Schrager-Lavelle for early assistance with *awl1* and brachypodium transformation; Chris Phillips and Dan Jones for greenhouse maintenance and plant growth; and Mary Byrne for providing us with *awl1* seed. We thank Amy Cartwright and John Vogel for assistance with sequencing. We also thank Kirstie Halsey and Hannah Walpole from Rothamsted Research’s Bioimaging team and staff of Rothamsted Research’s Horticulture and Controlled Environment Department. We thank Kirk Amundson, Hailong Yang, and critical friend Zachary Lippman for their thoughtful comments on the manuscript. This work was supported by the National Science Foundation (IOS-1652380, to MB), the Botanical Society of America’s Donald R. Kaplan Dissertation Award in Comparative Morphology (to ELP), a Lotta M. Crabtree Fellowship (to ELP), a USDA NIFA Postdoctoral Fellowship (2019-67012-29654) to JPG, and the UMass Natural History Collections. Rothamsted Research receives strategic funding from the Biotechnology and Biological Sciences Research Council of the United Kingdom (BBSRC) and DM acknowledges support from the Growing Health Institute Strategic Programme (BB/X010953/1; BBS/E/RH/230003A).

## Author contributions

EP, DM, and MB designed the research. EP, DM, MH, JG, DO, and BN performed experiments. EP analyzed data. EP and MB wrote the manuscript. All authors read and approved the final manuscript.

## Data availability

Short read sequencing data is available from the NCBI short read archive (PRJNA1074741). Data for ancestral state estimations are included in a dryad data archive (DOI: 10.5061/dryad.pc866t1zd).

## References

Abebe T, Wise RP, Skadsen RW. 2009. Comparative transcriptional profiling established the awn as the major photosynthetic organ of the barley spike while the lemma and the palea primarily protect the seed. The plant genome 2: 247–259.

Baulechner D, Jauker F, Neubauer TA, Wolters V. 2020. Convergent evolution of specialized generalists: Implications for phylogenetic and functional diversity of carabid feeding groups. Ecology and evolution 10: 11100–11110.

Blount ZD, Lenski RE, Losos JB. 2018. Contingency and determinism in evolution: Replaying life’s tape. Science 362.

Bolger AM, Lohse M, Usadel B. 2014. Trimmomatic: a flexible trimmer for Illumina sequence data. Bioinformatics 30: 2114–2120.

Bollback JP. 2006. SIMMAP: stochastic character mapping of discrete traits on phylogenies. BMC bioinformatics 7: 88.

Boyko JD, Beaulieu JM. 2021. Generalized hidden Markov models for phylogenetic comparative datasets. Methods in ecology and evolution / British Ecological Society 12: 468–478.

Cai L, Comont D, MacGregor D, Lowe C, Beffa R, Neve P, Saski C. 2023. The blackgrass genome reveals patterns of non-parallel evolution of polygenic herbicide resistance. The New phytologist 237: 1891–1907.

Cavanagh AM, Morgan JW, Godfree RC. 2020. Awn morphology influences dispersal, microsite selection and burial of Australian native grass diaspores. Frontiers in ecology and evolution 8.

Ceradini JP, Chalfoun AD. 2017. Species’ traits help predict small mammal responses to habitat homogenization by an invasive grass. Ecological applications: a publication of the Ecological Society of America 27: 1451–1465.

Clayton WD, Vorontsova MS, Harman KT, Williamson H. 2006 onwards. GrassBase - The Online World Grass Flora. http://www.kew.org/data/grasses-db.html.

Concordet J-P, Haeussler M. 2018. CRISPOR: intuitive guide selection for CRISPR/Cas9 genome editing experiments and screens. Nucleic acids research 46: W242–W245.

Derbyshire P, Byrne ME. 2013. MORE SPIKELETS1 is required for spikelet fate in the inflorescence of Brachypodium. Plant physiology 161: 1291–1302.

Duval-Jouve J. 1871. Étude anatomique de l’arête des graminées, par J. Duval-Jouve. J.-B. Baillière et fils.

Elbaum R, Zaltzman L, Burgert I, Fratzl P. 2007. The role of wheat awns in the seed dispersal unit. Science 316: 884–886.

Engler C, Youles M, Gruetzner R, Ehnert T-M, Werner S, Jones JDG, Patron NJ, Marillonnet S. 2014. A golden gate modular cloning toolbox for plants. ACS synthetic biology 3: 839–843.

Fourquin C, Primo A, Martínez-Fernández I, Huet-Trujillo E, Ferrándiz C. 2014. The CRC orthologue from Pisum sativum shows conserved functions in carpel morphogenesis and vascular development. Annals of botany 114: 1535–1544.

Fritz SA, Purvis A. 2010. Selectivity in mammalian extinction risk and threat types: a new measure of phylogenetic signal strength in binary traits. Conservation biology: the journal of the Society for Conservation Biology 24: 1042–1051.

Garnier LKM, Dajoz I. 2001. EVOLUTIONARY SIGNIFICANCE OF AWN LENGTH VARIATION IN A CLONAL GRASS OF FIRE-PRONE SAVANNAS. Ecology 82: 1720–1733.

Gould SJ. 2002. The Structure of Evolutionary Theory. Harvard University Press.

Grundbacher FJ. 1963. The physiological function of the cereal awn. The Botanical review; interpreting botanical progress 29: 366–381.

Grützner R, Martin P, Horn C, Mortensen S, Cram EJ, Lee-Parsons CWT, Stuttmann J, Marillonnet S. 2021. High-efficiency genome editing in plants mediated by a Cas9 gene containing multiple introns. Plant communications 2: 100135.

Harlan HV, Hulton HFE. 1920. Development of Barley Kernels in Normal and Clipped Spikes and the Limitations of Awnless and Hooded Varieties. Journal of the Institute of Brewing. Institute of Brewing 26: 639–641.

Hensen I, Müller C. 1997. Experimental and structural investigations of anemochorous dispersal. Plant Ecology 133: 169–180.

Hepler NK, Bowman A, Carey RE, Cosgrove DJ. 2020. Expansin gene loss is a common occurrence during adaptation to an aquatic environment. The Plant journal: for cell and molecular biology 101: 666–680.

Hong S-Y, Seo PJ, Yang M-S, Xiang F, Park C-M. 2008. Exploring valid reference genes for gene expression studies in Brachypodium distachyon by real-time PCR. BMC plant biology 8: 112.

Horn JW, van Ee BW, Morawetz JJ, Riina R, Steinmann VW, Berry PE, Wurdack KJ. 2012. Phylogenetics and the evolution of major structural characters in the giant genus Euphorbia L. (Euphorbiaceae). Molecular phylogenetics and evolution 63: 305–326.

Hsu H-C, Kuo Y-F. 2021. Nectar Guide Patterns on Developmentally Homologous Regions of the Subtribe Ligeriinae (Gesneriaceae). Frontiers in plant science 12: 650836.

Hua L, Wang DR, Tan L, Fu Y, Liu F, Xiao L, Zhu Z, Fu Q, Sun X, Gu P. 2015. LABA1, a domestication gene associated with long, barbed awns in wild rice. The Plant cell 27: 1875–1888.

Humphreys AM, Antonelli A, Pirie MD, Linder HP. 2011. ECOLOGY AND EVOLUTION OF THE DIASPORE ‘BURIAL SYNDROME’. Evolution; international journal of organic evolution 65: 1163–1180.

International Brachypodium Initiative. 2010. Genome sequencing and analysis of the model grass Brachypodium distachyon. Nature 463: 763–768.

James ME, Brodribb T, Wright IJ, Rieseberg LH, Ortiz-Barrientos D. 2023. Replicated Evolution in Plants. Annual review of plant biology 74: 697–725.

Jing Y, Wenbo C, Zhifeng H, Yan X, XinFang Z, Mi W, RuHui W, Wenqiang S, Jun Z, QianNan D, et al. 2023. DEGENERATED LEMMA (DEL) regulates lemma development and affects rice grain yield. Physiology and molecular biology of plants: an international journal of functional plant biology 29: 335–347.

Jin J, Hua L, Zhu Z, Tan L, Zhao X, Zhang W, Liu F, Fu Y, Cai H, Sun X, et al. 2016. GAD1 Encodes a Secreted Peptide That Regulates Grain Number, Grain Length and Awn Development in Rice Domestication. The Plant cell: tpc.00379.2016.

Kamilar JM, Cooper N. 2013. Phylogenetic signal in primate behaviour, ecology and life history. Philosophical transactions of the Royal Society of London. Series B, Biological sciences 368: 20120341.

Kaplan DR. 2001. Fundamental Concepts of Leaf Morphology and Morphogenesis: A Contribution to the Interpretation of Molecular Genetic Mutants. International journal of plant sciences 162: 465–474.

Kellogg EA. 2015. Flowering Plants. Monocots: Poaceae. Springer International Publishing.

Lee J-Y, Baum SF, Alvarez J, Patel A, Chitwood DH, Bowman JL. 2005. Activation of CRABS CLAW in the Nectaries and Carpels of Arabidopsis. The Plant cell 17: 25–36.

Lee W-S, Rudd JJ, Kanyuka K. 2015. Virus induced gene silencing (VIGS) for functional analysis of wheat genes involved in Zymoseptoria tritici susceptibility and resistance. Fungal genetics and biology: FG & B 79: 84–88.

Leichty AR, Sinha NR. 2021. A Grand Challenge in Development and Evodevo: Quantifying the Role of Development in Evolution. Frontiers in plant science 12: 752344.

Lewis MW, Hake S. 2016. Keep on growing: building and patterning leaves in the grasses. Current opinion in plant biology 29: 80–86.

Liller CB, Walla A, Boer MP, Hedley P, Macaulay M, Effgen S, von Korff M, van Esse GW, Koornneef M. 2017. Fine mapping of a major QTL for awn length in barley using a multiparent mapping population. TAG. Theoretical and applied genetics. Theoretische und angewandte Genetik 130: 269–281.

Linder HP, Lehmann CER, Archibald S, Osborne CP, Richardson DM. 2018. Global grass (Poaceae) success underpinned by traits facilitating colonization, persistence and habitat transformation. Biological reviews of the Cambridge Philosophical Society 93: 1125–1144.

Li X, Tang Y, Zhou C, Lv J. 2023. Contributions of glume and awn to photosynthesis, 14C assimilates and grain weight in wheat ears under drought stress. Heliyon 9: e21136.

Li X, Wang H, Li H, Zhang L, Teng N, Lin Q, Wang J, Kuang T, Li Z, Li B. 2006. Awns play a dominant role in carbohydrate production during the grain-filling stages in wheat (Triticum aestivum). Physiologia plantarum 127: 701–709.

Li B, Xu F. 2020. Homology and functions of inner staminodes in Anaxagorea javanica (Annonaceae). AoB plants 12: laa057.

Maydup ML, Antonietta M, Guiamet JJ, Graciano C, López JR, Tambussi EA. 2010. The contribution of ear photosynthesis to grain filling in bread wheat (Triticum aestivum L.). Field crops research 119: 48–58.

Ma X, Zhang J, Han B, Tang J, Cui D, Han L. 2019. FLA, which encodes a homolog of UBP, is required for chlorophyll accumulation and development of lemma and palea in rice. Plant cell reports 38: 321–331.

McAllister CA, McKain MR, Li M, Bookout B, Kellogg EA. 2018. Specimen-based analysis of morphology and the environment in ecologically dominant grasses: the power of the herbarium. Philosophical transactions of the Royal Society of London. Series B, Biological sciences 374.

McGee MD, Wainwright PC. 2013. Convergent evolution as a generator of phenotypic diversity in threespine stickleback. Evolution; international journal of organic evolution 67: 1204–1208.

Mellado-Sánchez M, McDiarmid F, Cardoso V, Kanyuka K, MacGregor DR. 2020. Virus-Mediated Transient Expression Techniques Enable Gene Function Studies in Black-Grass. Plant physiology 183: 455–459.

Miao J, Guo D, Zhang J, Huang Q, Qin G, Zhang X, Wan J, Gu H, Qu L-J. 2013. Targeted mutagenesis in rice using CRISPR-Cas system. Cell research 23: 1233–1236.

Miller EC, Gauch HG, Gries GA. 1944. A Study of the Morphological Nature and Physiological Functions of the Awns of Winter Wheat. Kansas State College of Agriculture and Applied Science Technical Bulletin 57.

Morris CD. 2021. Is a long hygroscopic awn an advantage for Themeda triandra in drier areas? African Journal of Range & Forage Science 38: 179–183.

Motzo R, Giunta F. 2002. Awnedness affects grain yield and kernel weight in near-isogenic lines of durum wheat. Australian journal of agricultural research 53: 1285–1293.

Muhsin M, Nawaz M, Khan I, Chattha MB, Khan S, Aslam MT, Iqbal MM, Amin MZ, Ayub MA, Anwar U, et al. 2021. Efficacy of Seed Size to Improve Field Performance of Wheat under Late Sowing Conditions. Pakistan Journal of Agricultural Research 34: 247–253.

Nagasawa N, Miyoshi M, Sano Y, Satoh H, Hirano H, Sakai H, Nagato Y. 2003. SUPERWOMAN1 and DROOPING LEAF genes control floral organ identity in rice. Development 130: 705–718.

Nik MM, Babaeian M, Tavassoli A. 2011. Effect of seed size and genotype on germination characteristic and seed nutrient content of wheat. Scientific Research and Essays 6: 2019–2025.

Ntakirutimana, Xie. 2019. Morphological and Genetic Mechanisms Underlying Awn Development in Monocotyledonous Grasses. Genes 10: 573.

Ntakirutimana F, Xie W. 2020. Unveiling the Actual Functions of Awns in Grasses: From Yield Potential to Quality Traits. International journal of molecular sciences 21.

O’Connor DL, Runions A, Sluis A, Bragg J, Vogel JP, Prusinkiewicz P, Hake S. 2014. A division in PIN-mediated auxin patterning during organ initiation in grasses. PLoS computational biology 10: e1003447.

Ohmori Y, Toriba T, Nakamura H, Ichikawa H, Hirano H-Y. 2011. Temporal and spatial regulation of DROOPING LEAF gene expression that promotes midrib formation in rice. The Plant journal: for cell and molecular biology 65: 77–86.

Orme D. 2013. The caper package: comparative analysis of phylogenetics and evolution in R.

Patterson EL, Richardson A, Bartlett M. 2023. Pushing the boundaries of organ identity: Homology of the grass lemma. American journal of botany 110: e16161.

Peart MH. 1979. Experiments on the Biological Significance of the Morphology of Seed-Dispersal Units in Grasses. The Journal of ecology 67: 843–863.

Peart MH. 1981. Further Experiments on the Biological Significance of the Morphology of Seed-Dispersal Units in Grasses. The Journal of ecology 69: 425–436.

Petersen KB, Kellogg EA. 2022. Diverse ecological functions and the convergent evolution of grass awns. American journal of botany 109: 1331–1345.

Peterson PM, Romaschenko K, Soreng RJ, Reyna JV. 2019. A key to the North American genera of Stipeae (Poaceae, Pooideae) with descriptions and taxonomic names for species of Eriocoma, Neotrinia, Oloptum, and five new genera: Barkworthia, ×Eriosella, Pseudoeriocoma, Ptilagrostiella, and Thorneochloa. PhytoKeys 126: 89–125.

Prostak SM, Robinson KA, Titus MA, Fritz-Laylin LK. 2021. The actin networks of chytrid fungi reveal evolutionary loss of cytoskeletal complexity in the fungal kingdom. Current biology: CB 31: 1192–1205.e6.

Prusinkiewicz P, Erasmus Y, Lane B, Harder LD, Coen E. 2007. Evolution and development of inflorescence architectures. Science 316: 1452–1456.

Rajakumar R, San Mauro D, Dijkstra MB, Huang MH, Wheeler DE, Hiou-Tim F, Khila A, Cournoyea M, Abouheif E. 2012. Ancestral developmental potential facilitates parallel evolution in ants. Science 335: 79–82.

Raju MVS. 2011. Studies on the inflorescence of wild oats (Avena fatua). Morphology and anatomy of the awn in relation to its movement. Canadian journal of botany. Journal canadien de botanique.

Ren D, Hu J, Xu Q, Cui Y, Zhang Y, Zhou T, Rao Y, Xue D, Zeng D, Zhang G, et al. 2018. FZP determines grain size and sterile lemma fate in rice. Journal of experimental botany 69: 4853–4866.

Revell LJ. 2012. phytools: an R package for phylogenetic comparative biology (and other things). Methods in ecology and evolution / British Ecological Society 3: 217–223.

Sajo MG, Longhi-Wagner HM, Rudall PJ. 2008. Reproductive morphology of the early-divergent grass Streptochaeta and its bearing on the homologies of the grass spikelet. Plant systematics and evolution = Entwicklungsgeschichte und Systematik der Pflanzen 275: 245–255.

Sanchez-Bragado R, Molero G, Araus JL, Slafer GA. 2023. Awned versus awnless wheat spikes: does it matter? Trends in plant science 28: 330–343.

Schöning C, Espadaler X, Hensen I, Roces F. 2004. Seed predation of the tussock-grass Stipa tenacissima L. by ants (Messor spp.) in south-eastern Spain: the adaptive value of trypanocarpy. Journal of arid environments 56: 43–61.

Schubert M, Marcussen T, Meseguer AS, Fjellheim S. 2019. The grass subfamily Pooideae: Cretaceous–Palaeocene origin and climate-driven Cenozoic diversification. Global ecology and biogeography: a journal of macroecology 28: 1168–1182.

Shoesmith JR, Solomon CU, Yang X, Wilkinson LG, Sheldrick S, van Eijden E, Couwenberg S, Pugh LM, Eskan M, Stephens J, et al. 2021. APETALA2 functions as a temporal factor together with BLADE-ON-PETIOLE2 and MADS29 to control flower and grain development in barley. Development 148.

Sifuentes-Romero I, Ferrufino E, Thakur S, Laboissonniere LA, Solomon M, Smith CL, Keene AC, Trimarchi JM, Kowalko JE. 2020. Repeated evolution of eye loss in Mexican cavefish: Evidence of similar developmental mechanisms in independently evolved populations. Journal of experimental zoology. Part B, Molecular and developmental evolution 334: 423–437.

Song Y-X, Peng S, Mutie FM, Jiang H, Ren J, Cong Y-Y, Hu G-W. 2022. Evolution and Taxonomic Significance of Seed Micromorphology in Impatiens (Balsaminaceae). Frontiers in plant science 13: 835943.

Strable J, Nelissen H. 2021. The dynamics of maize leaf development: Patterned to grow while growing a pattern. Current opinion in plant biology 63: 102038.

Strable J, Vollbrecht E. 2019. Maize YABBY genes drooping leaf1 and drooping leaf2 regulate floret development and floral meristem determinacy. Development 146: dev171181.

Strable J, Wallace JG, Unger-Wallace E, Briggs S, Bradbury PJ, Buckler ES, Vollbrecht E. 2017. Maize YABBY Genes drooping leaf1 and drooping leaf2 Regulate Plant Architecture. The Plant cell 29: 1622–1641.

Swarts K, Gutaker RM, Benz B, Blake M, Bukowski R, Holland J, Kruse-Peeples M, Lepak N, Prim L, Romay MC, et al. 2017. Genomic estimation of complex traits reveals ancient maize adaptation to temperate North America. Science 357: 512–515.

Teisher JK, McKain MR, Schaal BA, Kellogg EA. 2017. Polyphyly of Arundinoideae (Poaceae) and evolution of the twisted geniculate lemma awn. Annals of botany 120: 725–738.

Thi-Tuyet-Hoa MT. 1965. Les glumelles inférieures aristées de quelques Graminées: anatomie, morphologie. Bulletin du Jardin botanique de l’État a Bruxelles 35: 219–284.

Thorvaldsdóttir H, Robinson JT, Mesirov JP. 2013. Integrative Genomics Viewer (IGV): high-performance genomics data visualization and exploration. Briefings in bioinformatics 14: 178–192.

Titulaer M, Melgoza-Castillo A, Macías-Duarte A, Panjabi AO. 2018. Seed size, bill morphology, and handling time influence preferences for native vs. nonnative grass seeds in three declining sparrows. The Wilson Journal of Ornithology 130: 445–456.

Toriba T, Hirano H-Y. 2014. The DROOPING LEAF and OsETTIN2 genes promote awn development in rice. The Plant journal: for cell and molecular biology 77: 616–626.

Torruella G, de Mendoza A, Grau-Bové X, Antó M, Chaplin MA, del Campo J, Eme L, Pérez-Cordón G, Whipps CM, Nichols KM, et al. 2015. Phylogenomics Reveals Convergent Evolution of Lifestyles in Close Relatives of Animals and Fungi. Current biology: CB 25: 2404–2410.

Vogel J, Hill T. 2008. High-efficiency Agrobacterium-mediated transformation of Brachypodium distachyon inbred line Bd21-3. Plant cell reports 27: 471–478.

Von Goethe JW, Miller GL. 2009. The metamorphosis of plants. MIT Press Cambridge.

Ware A, Jones DH, Flis P, Chrysanthou E, Smith KE, Kümpers BMC, Yant L, Atkinson JA, Wells DM, Bhosale R, et al. 2023. Loss of ancestral function in duckweed roots is accompanied by progressive anatomical reduction and a re-distribution of nutrient transporters. Current biology: CB 33: 1795–1802.e4.

Webster MA, Gilmartin PM. 2006. Analysis of late stage flower development in Primula vulgaris reveals novel differences in cell morphology and temporal aspects of floral heteromorphy. The New phytologist 171: 591–603.

Wessinger CA, Hileman LC. 2016. Accessibility, constraint, and repetition in adaptive floral evolution. Developmental biology 419: 175–183.

West-Eberhard MJ. 2003. Plasticity. In: Oxford University Press.

Weyhrich RA, Carver BF, Smith EL. 1994. Effect of awn suppression on grain yield and agronomic traits in hard red winter wheat. Crop science 34: 965–969.

Wu TD, Reeder J, Lawrence M, Becker G, Brauer MJ. 2016. GMAP and GSNAP for Genomic Sequence Alignment: Enhancements to Speed, Accuracy, and Functionality. Methods in molecular biology 1418: 283–334.

Yamaguchi T, Nagasawa N, Kawasaki S, Matsuoka M, Nagato Y, Hirano H-Y. 2004. The YABBY gene DROOPING LEAF regulates carpel specification and midrib development in Oryza sativa. The Plant Cell Online 16: 500–509.

Yanez A, Desta I, Commins P, Magzoub M, Naumov P. 2018. Morphokinematics of the Hygroactuation of Feather Grass Awns. Advanced Biosystems 2: 1800007.

Yuo T, Yamashita Y, Kanamori H, Matsumoto T, Lundqvist U, Sato K, Ichii M, Jobling SA, Taketa S. 2012. A SHORT INTERNODES (SHI) family transcription factor gene regulates awn elongation and pistil morphology in barley. Journal of experimental botany 63: 5223–5232.

Zhang Y, Shen C, Li G, Shi J, Yuan Y, Ye L, Song Q, Shi J, Zhang D. 2024. MADS1-regulated lemma and awn development benefits barley yield. Nature communications 15: 301.

Zhang Y, Yu H, Liu J, Wang W, Sun J, Gao Q, Zhang Y, Ma D, Wang J, Xu Z, et al. 2016. Loss of function of OsMADS34 leads to large sterile lemma and low grain yield in rice (Oryza sativa L.). Molecular breeding: new strategies in plant improvement 36: 147.

Zhang L, Zhu X, Zhao Y, Guo J, Zhang T, Huang W, Huang J, Hu Y, Huang C-H, Ma H. 2022. Phylotranscriptomics Resolves the Phylogeny of Pooideae and Uncovers Factors for Their Adaptive Evolution. Molecular biology and evolution 39.

