## Supplemental Figures and Tables for "Deep Homology and Developmental Constraint Underlies the Replicated Evolution of Grass Awns"

**Article title:** Deep homology and developmental constraint underlie the replicated evolution of grass awns

The following supporting information is available for this article:

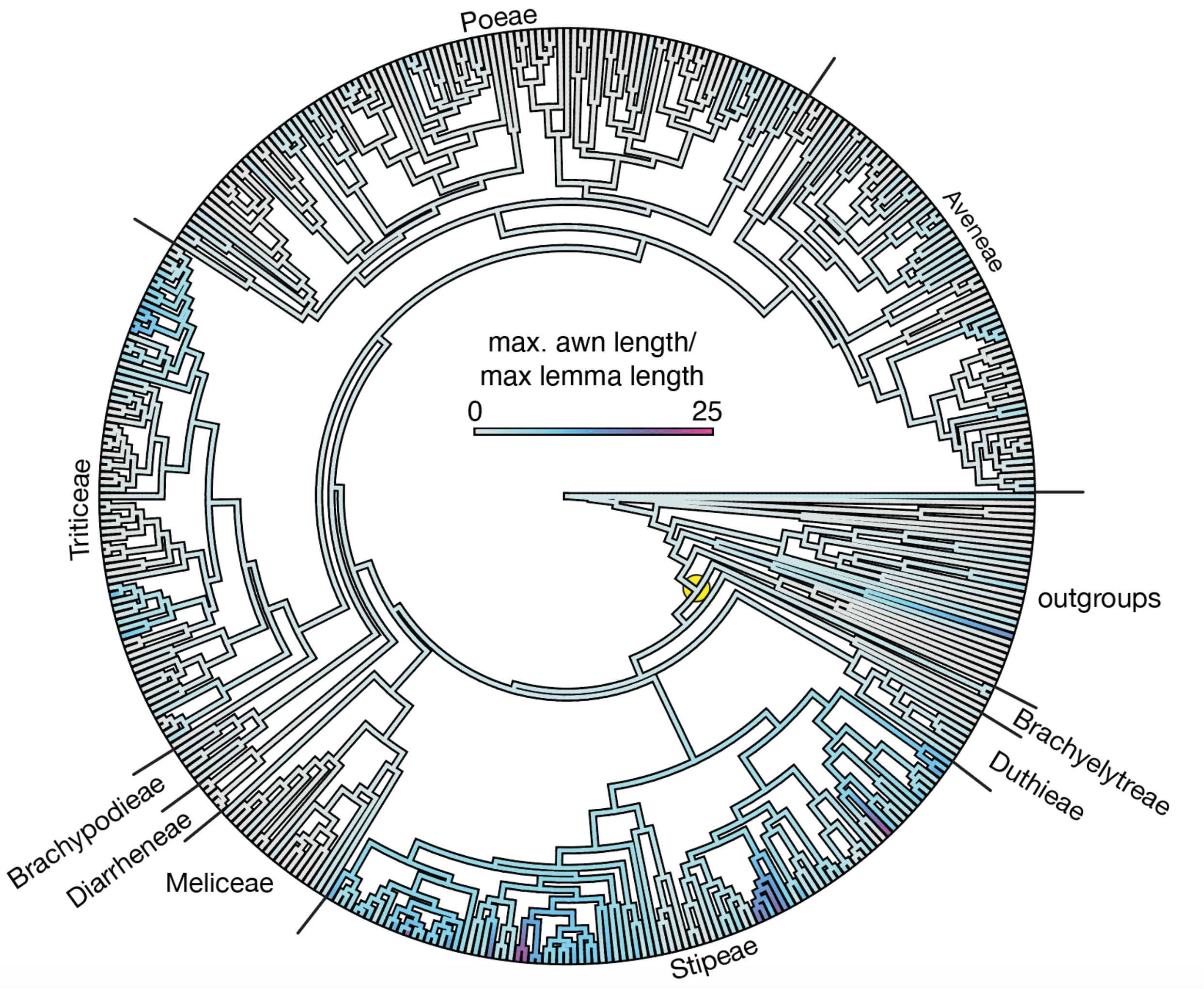


**Fig. S1.** Ancestral state reconstruction of (awn length/lemma length), focused on the Pooideae.

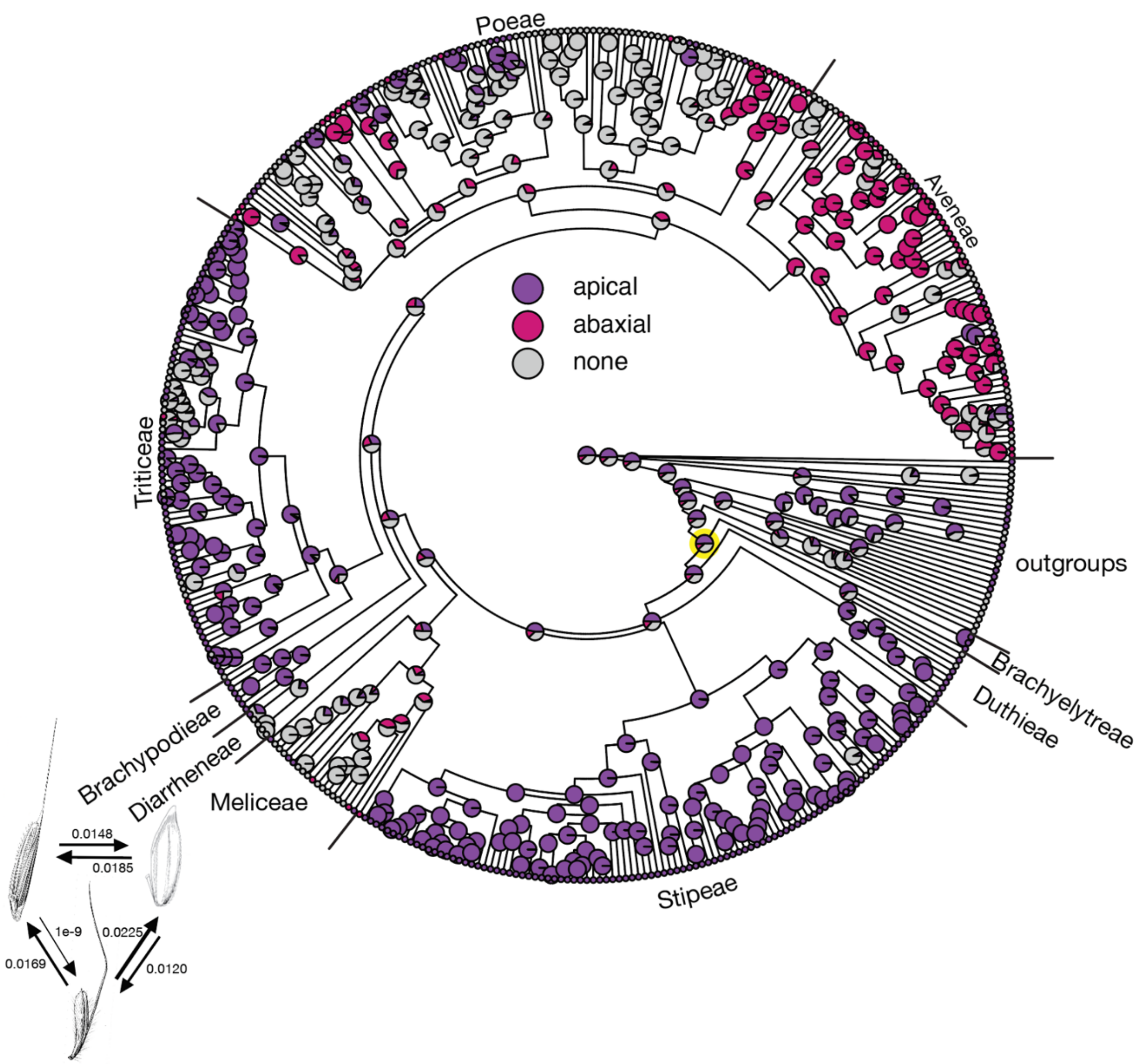


**Fig. S2.** Ancestral state reconstruction for awn insertion point, focused on the Pooideae. Yellow node: most recent common ancestor. Transition rates between states shown in bottom left.

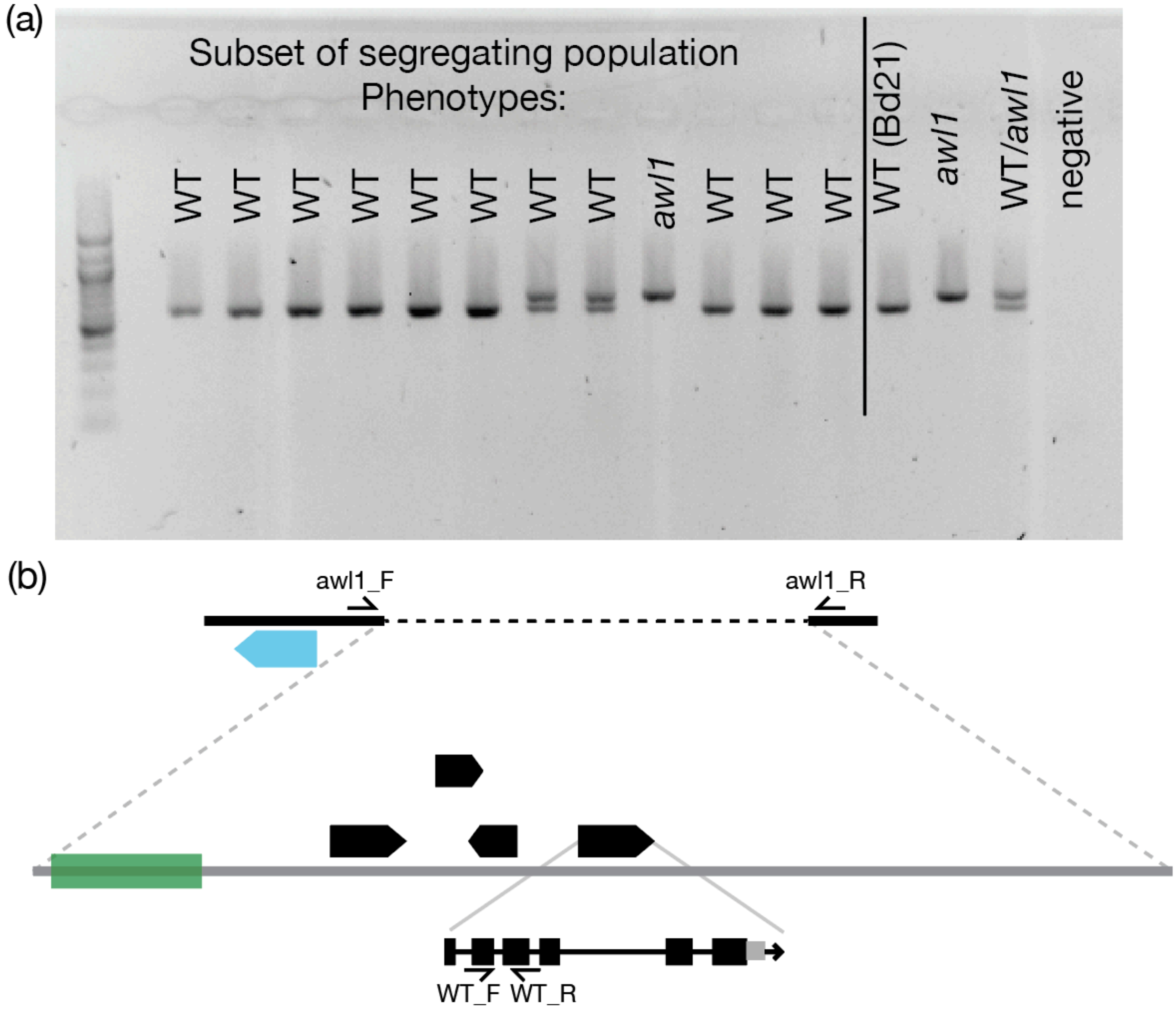


**Fig. S3.** *awl1* segregates as a single-locus mutation. (a) Gel electrophoresis of PCR assay for *awl1* deletion. (b) Diagram of primers for *awl1* deletion assay with multiplexed primers. Primers on either end of deletion (unable to amplify in WT), and primers within gene in deleted region (unable to amplify from *awl1*).

(a)

| | plants with awnless flowers |
|---|---|
| BSMV:GFP | 0/8 |
| BSMV: MCS | 0/8 |
| BSMV: PDS | 0/8 |

(b)

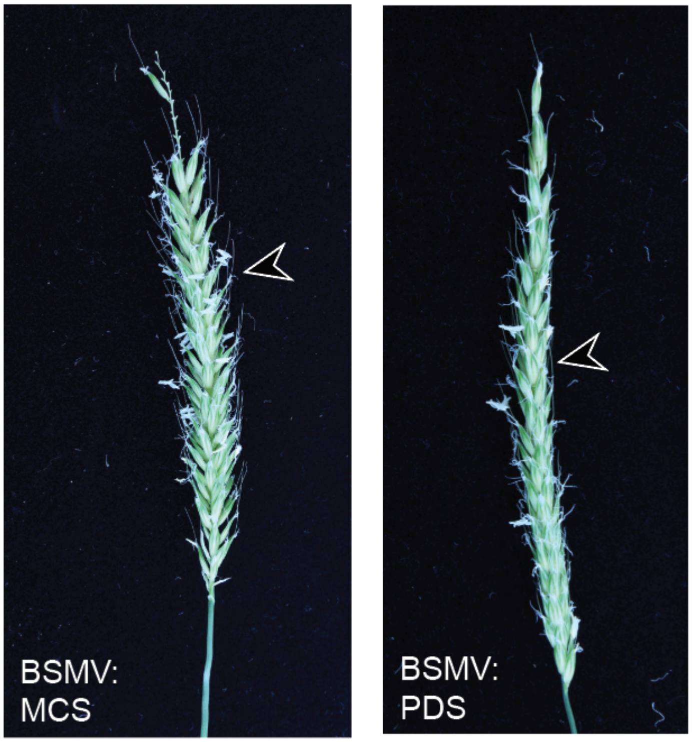


**Fig. S4.** Phenotypes for Multiple Cloning Site (empty vector) and phytoene desaturase (PDS) negative controls in VIGS experiment. (a) No control plants produced any awnless florets. (b) Typical spikes from the MCS and PDS controls, arrows note awns.

**Table S1.** Primers used in this study.

| Target | Oligo Type | Sequence | Application |
|---|---|---|---|
| **BdDL FW** | qPCR Primer | TTCTTCTCCGGGGGCTTCA | Gene expression (DL) |
| **BdDL RV** | qPCR Primer | CGTTCCAGGGATGCACTGA | |
| **BdUBC18 FW** | qPCR Primer | GGAGGCACCTCAGGTCATTT | Gene expression (control) |
| **BdUBC18 RV** | qPCR Primer | ATAGCGGTCATTGTCTTGCG | |
| **BdDL-CR2 FW** | PCR Primer | GCCACGCAACACAAAGGAAT | CRISPR-Cas9 editing genotyping (exon 2) |
| **BdDL-CR2 RV** | PCR Primer | CTCCGGGGCTACAACCAAAT | |
| **BdDL-CR4 FW** | PCR Primer | GCTCCCTTTGTTGTGAAGCG | CRISPR-Cas9 editing genotyping (exon 4) |
| **BdDL-CR4 RV** | PCR Primer | GGCTAAGTTGTGACGAGGGA | |
| ***awl1* deletion border FW** | PCR Primer | CACCTTTTGAAGCCCCATGT | *awl1* genotyping for gel assay in multiplex |
| ***awl1* deletion border RV** | PCR Primer | ACCTTGTGGACACCGGCTT | |
| ***awl1* deleted region FW** | PCR Primer | GTGTAACTGTGCAAACGAGC | *awl1* genotyping for gel assay in multiplex |
| ***awl1* deleted region RV** | PCR Primer | ATCTCTATCCCCGTCCTTTG | |
| **BdDL-CR exon 2** | CRISPR-Cas9 spacer | ACCTCTCCTTCCTCAGCCCG | CRISPR guides (DL) |
| **BdDL-CR exon 4** | CRISPR-Cas9 spacer | GATTGTAAGCAGATGGAAGG | |
